# Genetic diversity of the spinach downy mildew pathogen based on hierarchical sampling

**DOI:** 10.1101/2020.02.18.953661

**Authors:** Chunda Feng, Kurt Lamour, Braham Deep Singh Dhillon, Maria Isabel Villarroel-Zeballos, Vanina Lilian Castroagudin, Burt H. Bluhm, Ainong Shi, Alejandro Rojas, James C. Correll

## Abstract

Downy mildew, caused by the obligate oomycete pathogen *Peronospora effusa*, is the most economically important disease of spinach. In the past 30 years, 14 new races and 13 strains with novel virulence have been identified. However, the mechanism(s) driving the rapid evolution of virulence remains unknown. To understand reproductive strategies potentially driving the emergence of new races in *P. effusa*, 67 composite isolates (a collection of symptomatic leaves from a single cultivar grown in a defined area) of *P. effusa* obtained from 13 states between 2010 and 2018 were used to analyze the population genetic diversity hierarchically. Genotypes at 33 SNP loci of 719 lesions from these 67 isolates were determined by targeted sequencing. Diversity was then evaluated among individual lesions within the composite isolates, between isolates, host cultivars, geographic locations, and years of isolates collected. A total of 380 genotypes were identified from 719 individual lesions. Of the 380 genotypes, 350 (92%) were unique while the most common genotype was identified in 110 lesions of 16 isolates collected from 13 cultivars from CA and AZ in 2016. Variation within composite isolates ranged from none (a single genotype among lesions from a composite isolate) to 38 unique genotypes recovered from 39 lesions of a composite isolate. An index of association analysis suggested asexual (clonal) and sexual reproduction play important roles in population structure. Based on discriminant analysis of principal components, four distinct subpopulations were identified. Host cultivar, origin, and time of collection had an effect on population differentiation, and genotypes specific to a certain location or collection period were identified. Some subpopulations were unique to certain areas, and were only detected after 2014-2016. The co-existence of sexual and asexual reproduction strategies may partially explain the rapid emergence and spread of new races and novel strains of *P. effusa*.

## Introduction

Downy mildew, caused by the obligate oomycete pathogen *Peronospora effusa* (= *P. farinosa* f. sp. *spinaciae*), is the most economically important disease of spinach [1–3]. High downy mildew disease pressure can make spinach fields un-harvestable, and even low levels of disease may require culling infected leaves before harvest, thus decreasing profitability. In conventional spinach production, downy mildew is managed with resistant cultivars and/or fungicides. However, approximately 50% of U.S. spinach production is organic, which negates chemical control and makes host resistance the most practical management tool. Unfortunately, the pathogen evolves rapidly, For example, 14 of the 17 named races and 13 novel virulence strains of *P. effusa* have been identified in the past 30 years [4–8]. New races and novel strains of the pathogen may overcome newly released resistant cultivars, and result in disease epidemics and severe economic losses.

Despite the economic impact of spinach downy mildew, the mechanisms underlying rapid emergence of new races in *P. effusa* are unknown. One hypothesis is that mutations and/or epigenetic changes drive the emergence of new races. Modern practices of high-density, year-round spinach production are highly conducive for downy mildew and provide “green-bridges” that sustain pathogen survival. The pathogen produces copious numbers of asexual sporangia, and resistant cultivars exert strong selection pressure on the pathogen. Plant resistance genes directly or indirectly recognize corresponding pathogen avirulence genes, and trigger host immune responses [9]. Thus, the absence or functional disruption of an avirulence gene in the pathogen could enable infection on a previously resistant cultivar, which would give rise to a new race. Consistent with this hypothesis, the spinach cultivar Pigeon is resistant to race 12 but susceptible to race 14, and isolates of these two races have similar pathotypes on the panel of differential cultivars previously used for race determination [5]. When comparing 100 contigs of whole-genome sequences from UA2209 (race 12) and UA4410 (race 14), both isolates were highly heterozygous and the distribution of heterozygous sites were identical across 83 of the 100 contigs. However, isolate UA4410 (race 14) showed a loss of heterozygosity across 17 of the 100 contigs relative to UA2209 [10], which suggested the pathogen has a mechanism to alter, or “lose function” of specific alleles by altering zygosity. Thus, one possible inference is that races could evolve by losing the function of one or more avirulence alleles through a phenomenon known as “loss of heterozygosity” [11]. Another hypothesis is that the pathogen reshuffles virulence/avirulence genes during sexual reproduction, resulting in new races and/or deviating strains. Oospores of *P. effusa* have often been observed on spinach seeds [12–14]. Although oospores were historically thought to be rare on vegetative plant tissues [5], they have recently become substantially more common, which were found in two isolates of 2016 and 28 isolates of 2018 (Correll et al., unpublished). However, a study indicating that oospores from infested or infected seed can give rise to infected seedlings [13] has been difficult to reproduce despite robust efforts to confirm the phenomenon (Correll et al, unpublished). Thus, the frequency and importance of sexual reproduction as a driver of race emergence under field conditions has been difficult to establish. The seedborne transmission [12, 13, 15] and global distribution of some races of *P. effusa* [5] suggested that new races may be introduced to other spinach production regions.

The emergence and/or introduction of new races and genotypes would alter the genetic diversity of the *P. effusa* population. Tremendous variation has been observed in many other plant-pathogenic oomycetes [16–19]. A number of mechanisms may contribute to the genetic variability of these oomycetes, including 1) during asexual reproduction, mitotic recombination and transposable elements could result in genetic variation [19, 20]; 2) The meiotic recombination due to sexual reproduction could result in significant variation in the progeny and better adaptation to the environment [19, 21–23]. Plant pathogenic oomycetes can secrete effectors to manipulate host resistance systems and initiate infection [24–26]. Hundreds of effectors have been identified in oomycetes [27], occupying chromosome regions rich in repetitive DNA that might lead to accelate effector evolution by non-allelic homologous recombination and tandem gene duplication [28]. The arsenal of combined effectors from both parents as a result of sexual reproduction enable the pathogen to infect a broader range of host cultivars; 3) In both sexual and asexual reproduction, the repairing of mismatchbases during homologous recombination may result in gene conversion, a process where one DNA sequence replaces its homologous sequence so that the sequences become identical after the conversion [19, 20, 29]; 4) Heterokaryosis by the fusion of hyphae in fungi and oomycetes could lead to genetic variation such as aneuploidy [30–34]. However, the genetic diversity of *P. effusa* populations has not been investigated thoroughly. Historically, efforts have focused on studying the phenotypic diversity of isolates of *P. effusa*, with emphasis on determining pathogenicity on differential spinach cultivars (e.g., determining race). An initial exploration of genotypic diversity in populations of *P. effusa* has been conducted with samples collected during 2008-2015 from CA and AZ [10]. However, little is known about the genetic diversity of *P. effusa* within a composite isolate, from a given host cultivar, or the spatial and temporal impacts on the genetic diversity on the pathogen.

The objective of this research was to investigate the genetic diversity of *P. effusa* at different hierarchical levels, including within and among spinach downy mildew composite isolates collected from 13 states over an eight-year sampling period.

## Materials and Methods

### Spinach downy mildew isolates

The terminology used for a spinach downy mildew isolate is somewhat unconventional. For example, an isolate collected for race typing by the International Working Group on Perononspora (IWGP) on spinach, with Plantum (based in Gouda, Netherlands), involves collecting multiple infected leaves within a limited area in a field from a given cultivar at a given time. Getting an infection from a single spore culture, often used for fungal species, has proven to be problematic with this pathogen, as well as other obligate plant pathogens. Thus, for the purposes of this effort, the term composite isolate is used which is defined as a collection of individual infected leaves within a 100 square meter area, all collected from a single cultivar at a given time. A total of 67 spinach downy mildew composite isolates, including 62 samples collected from 2015 to 2018, and five samples collected from 2010 to 2014 (UA0510C, US1612B, UA4712, UA2213, and UA1014APLP), were evaluated in this study (Table 1).

**Table 1.**
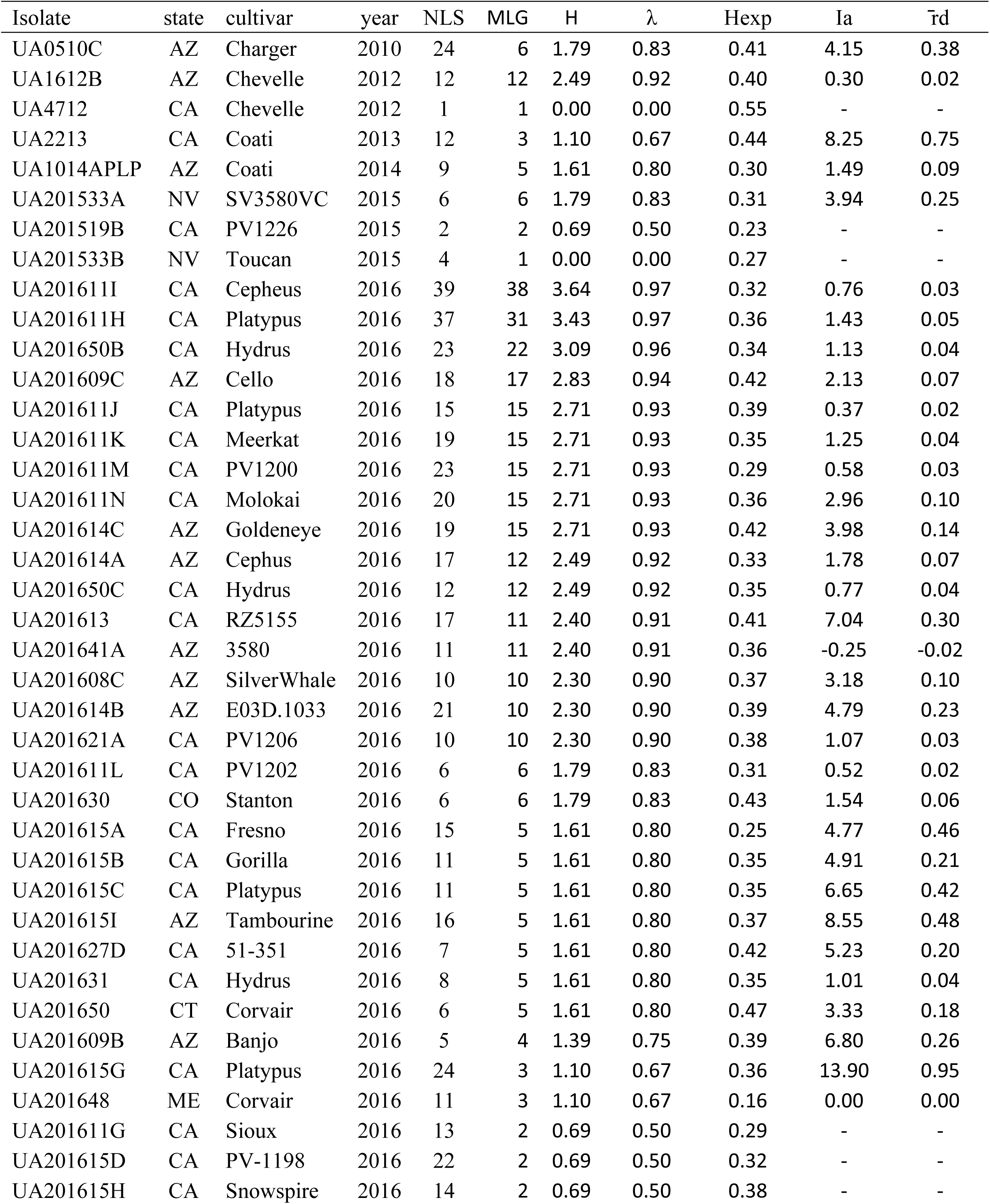
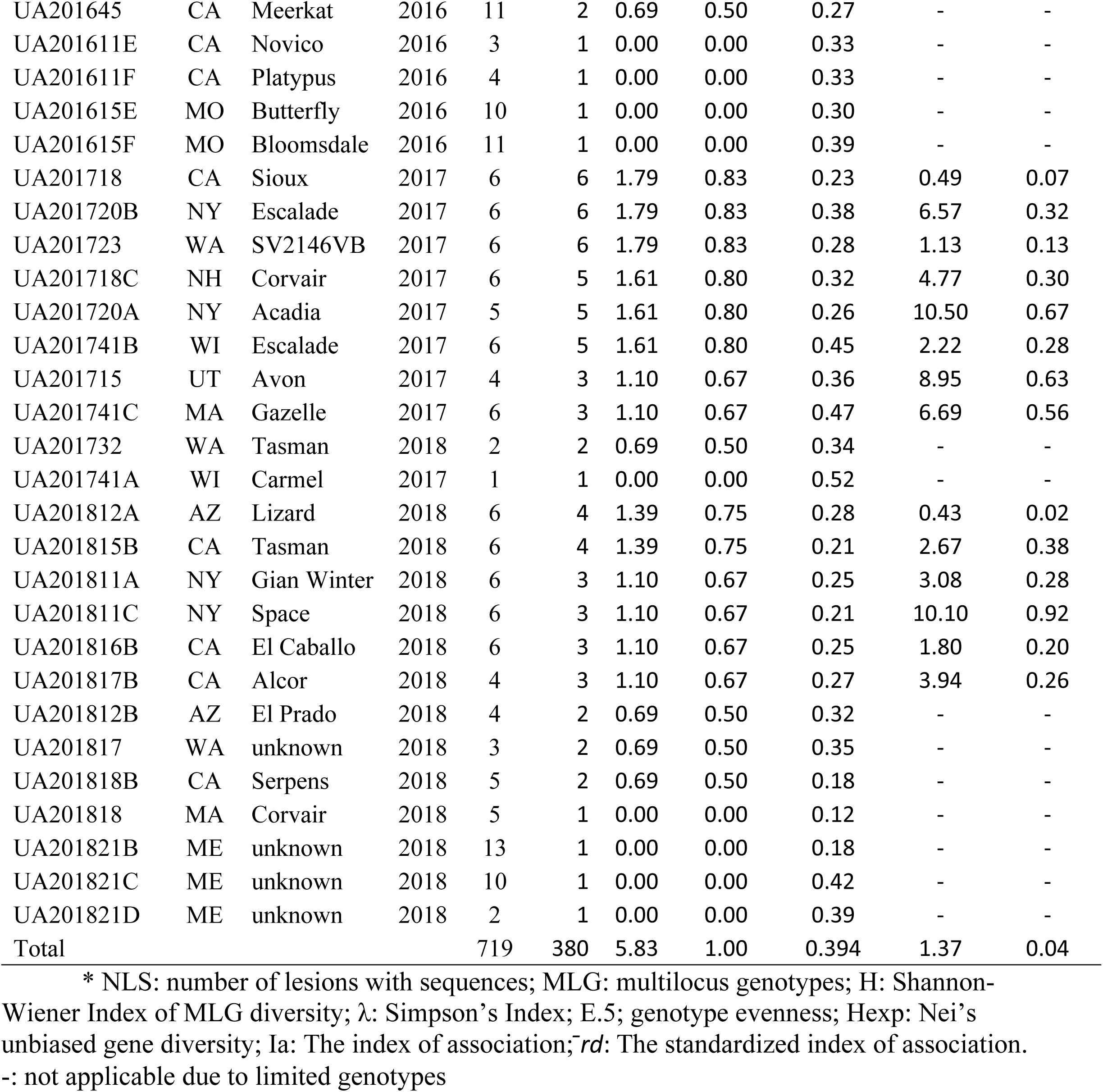
Spinach downy mildew isolates and diversity parameters after clone correction within each isolate*

### Genotyping with the targeted sequencing approach

For a finer-scale genetic analysis of the leaves comprising the composite isolate, targeted-sequencing was utilized to genotype individual lesions from individual leaves, the procedures were shown in Figure 1. Briefly, from each lesion, four leaf discs (approximately 7 mm diameter) were punched with a plastic straw and placed in one well of a 96-well plate, six to 40 lesions were processed for each isolate. Leaf tissue in the deep-well plates were lyophilized before DNA extraction using a silica based approach previously described [35]. A total of 33 SNP markers were selected and amplified in multiplex PCR reactions for each sample at Floodlight Genomics (Knoxville, TN) as previously described [10]. The PCR products in each well were subjected to targeted sequencing with an Illumina Hiseq-2000 system [10]. Thus, between 2 and 39 lesions with targeted-sequencing genotypes were examined for 65 composite isolates, and two composite isolates had only a lesion with single sequencing reaction each. SNP genotypes were assigned to target sites with at least 10x coverage.

**Fig. 1.**
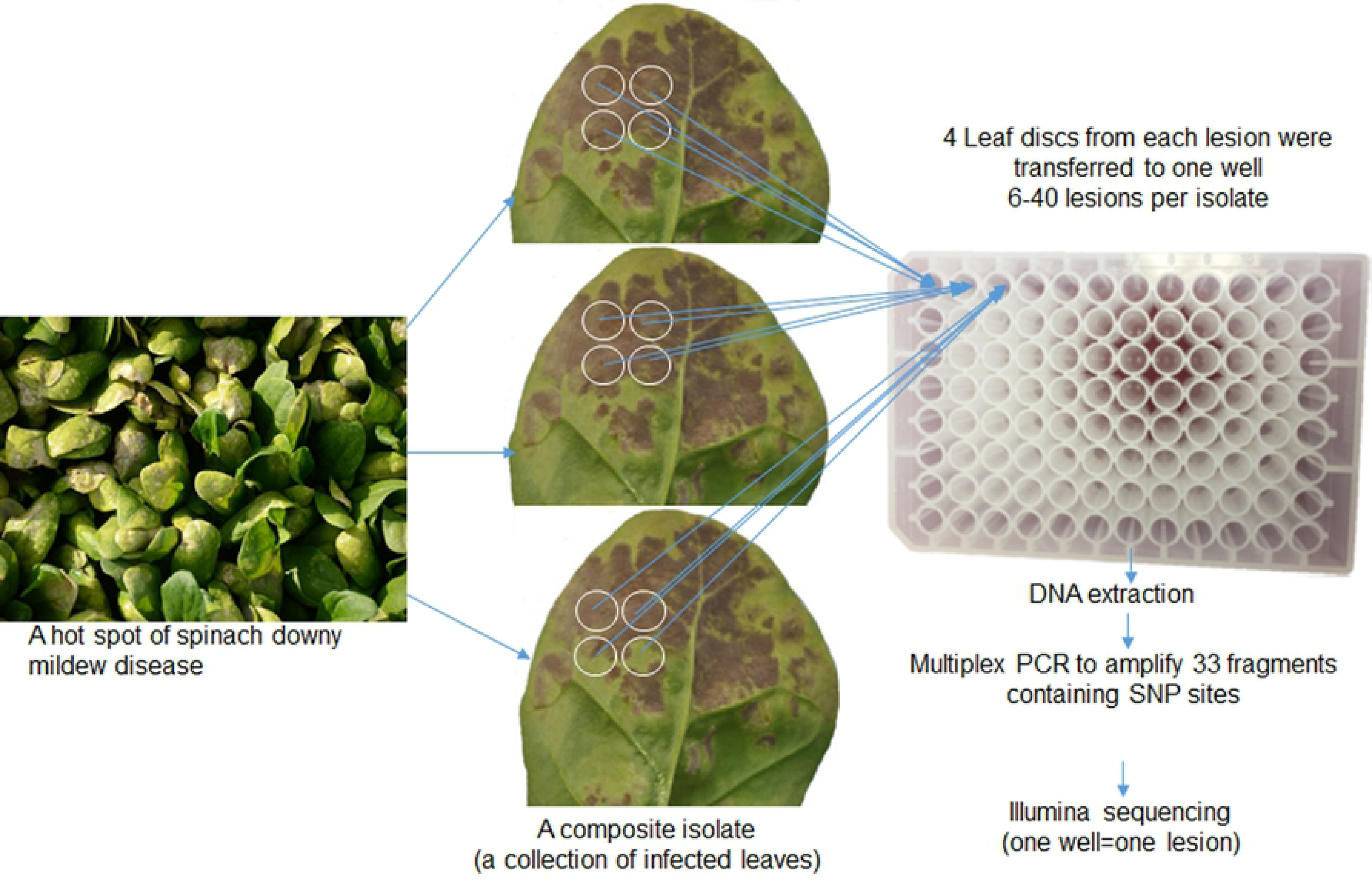
Schematic showing the procedures from collecting a spinach downy mildew composite isolate to genotyping lesions by targeted sequencing

### Data analysis

The population structure of *P. effusa* was analyzed at different hierarchical levels with the statistical language R using various R packages, including poppr [36, 37], adegenet [38, 39], and hierfstat [40], etc. Each of the 65 isolates with two or more lesions was analyzed independently to assess genetic variation within that isolate. Then, all lesions were analyzed according to the state origin, year, and host cultivar that the isolates were collected using stratification of year/state/cultivar/isolate hierarchical structure with and without clone correction. Clone correction was performed to remove duplicated genotypes at different level of strata after which the genetic diversity was assessed. An analysis of molecular variance (AMOVA) was performed to determine the impact of each hierarchary level on the genetic diversity of *P. effusa* [36]. A discriminant analysis of principal components (DAPC) was conducted to detect the clustering of the pathogen at different levels [41]. The index of association and the stantdardized index of association were tested for difference from zero by 1000 permutations, with zero an indication of linkage equilibrium from a sexually reproduced population, and significant greater than zero an indication of linkage disequilibrium from a clonal population or mixed reproduction [16, 36, 37].

## Results

### Genotypic and allelic analysis of spinach downy mildew isolates

Sequencing results were obtained from a total of 719 lesions from 67 composite isolates, including one lesion from each of isolates UA4712 and UA201741A, and 2 to 39 lesions from other 65 isolates (Table 1). Each lesion was genotyped at 33 SNP loci without missing data.

Allelic information of the 33 SNPs loci was listed in Table 2. Each of the 33 loci sequenced showed two alleles, which indicated *P. effusa* was diploid. There were two genotypes at each of three loci, and three genotypes at other loci. Major allele frequencies at each locus ranged from 0.53 to 0.95, with 5 greater than 0.9 (Table 2). Twenty-three loci were not in Hardy-Weinberg equilibrium (p<0.05) among all lesions. For each locus, the observe heterozygosity is larger than the heterozygosity of the total population, and both were greater than the heterozygosity of withing subdivided populations based on origin. The fixation index F_ST_ ranged from 0.02 to 0.26, with overall 0.13. All the inbreeding coefficient F_IS_ were negative, indicating that individuals in a population are less related (Table 2). All these results suggested population differentiation at most of the loci.

**Table 2.** Allelic information at 33 loci of 67 isolates of *P. effusa*\*

| locus | allele | genotype | MAF | Ho | Hs | Ht | Fst | Fis |
| --- | --- | --- | --- | --- | --- | --- | --- | --- |
| X1027_12305 | 2 | 3 | 0.66 | 0.63 | 0.39 | 0.44 | 0.12 | -0.64 |
| X1054_12307 | 2 | 3 | 0.55 | 0.44 | 0.35 | 0.44 | 0.20 | -0.25 |
| X1224_36034 | 2 | 3 | 0.85 | 0.54 | 0.35 | 0.40 | 0.10 | -0.53 |
| X14_20241 | 2 | 3 | 0.67 | 0.63 | 0.45 | 0.48 | 0.06 | -0.40 |
| X150_21870 | 2 | 3 | 0.76 | 0.54 | 0.39 | 0.48 | 0.20 | -0.39 |
| X157_954 | 2 | 3 | 0.91 | 0.04 | 0.05 | 0.05 | 0.04 | 0.04 |
| X162_20472 | 2 | 3 | 0.68 | 0.75 | 0.46 | 0.48 | 0.05 | -0.64 |
| X194_1933 | 2 | 3 | 0.6 | 0.47 | 0.38 | 0.48 | 0.21 | -0.23 |
| X230_11674 | 2 | 3 | 0.69 | 0.56 | 0.42 | 0.47 | 0.11 | -0.33 |
| X233_2053 | 2 | 3 | 0.62 | 0.46 | 0.39 | 0.47 | 0.17 | -0.18 |
| X234_4106 | 2 | 3 | 0.66 | 0.63 | 0.41 | 0.45 | 0.09 | -0.54 |
| X247_11808 | 2 | 3 | 0.77 | 0.45 | 0.30 | 0.36 | 0.15 | -0.47 |
| X288_15713 | 2 | 2 | 0.63 | 0.59 | 0.35 | 0.42 | 0.17 | -0.71 |
| X405_1475 | 2 | 3 | 0.67 | 0.31 | 0.30 | 0.39 | 0.24 | -0.04 |
| X425_3047 | 2 | 3 | 0.53 | 0.76 | 0.46 | 0.49 | 0.05 | -0.64 |
| X445_18260 | 2 | 3 | 0.91 | 0.05 | 0.05 | 0.05 | 0.03 | 0.03 |
| X505_1292 | 2 | 3 | 0.55 | 0.69 | 0.47 | 0.50 | 0.06 | -0.48 |
| X516_687 | 2 | 3 | 0.53 | 0.32 | 0.29 | 0.37 | 0.20 | -0.10 |
| X536_39392 | 2 | 3 | 0.67 | 0.52 | 0.34 | 0.39 | 0.13 | -0.52 |
| X597_1412 | 2 | 3 | 0.66 | 0.25 | 0.21 | 0.25 | 0.15 | -0.18 |
| X6_18952 | 2 | 3 | 0.56 | 0.77 | 0.48 | 0.50 | 0.04 | -0.60 |
| X606_18824 | 2 | 2 | 0.93 | 0.16 | 0.11 | 0.15 | 0.23 | -0.41 |
| X635_20522 | 2 | 3 | 0.59 | 0.51 | 0.36 | 0.43 | 0.15 | -0.41 |
| X638_43925 | 2 | 3 | 0.75 | 0.22 | 0.20 | 0.22 | 0.07 | -0.08 |
| X692_12307 | 2 | 3 | 0.84 | 0.51 | 0.30 | 0.38 | 0.20 | -0.67 |
| X703_46170 | 2 | 2 | 0.95 | 0.32 | 0.19 | 0.27 | 0.29 | -0.68 |
| X712_2077 | 2 | 3 | 0.73 | 0.53 | 0.33 | 0.39 | 0.16 | -0.59 |
| X76_6417 | 2 | 3 | 0.7 | 0.69 | 0.44 | 0.47 | 0.06 | -0.57 |
| X793_2335 | 2 | 3 | 0.69 | 0.69 | 0.43 | 0.49 | 0.11 | -0.59 |
| X795_9577 | 2 | 3 | 0.74 | 0.46 | 0.31 | 0.46 | 0.32 | -0.47 |
| X851_9386 | 2 | 3 | 0.66 | 0.47 | 0.32 | 0.38 | 0.15 | -0.44 |
| X863_51377 | 2 | 3 | 0.72 | 0.54 | 0.33 | 0.39 | 0.17 | -0.64 |
| X87_6382 | 2 | 3 | 0.93 | 0.39 | 0.22 | 0.31 | 0.29 | -0.74 |
| mean | 2 | 2.91 | 0.71 | 0.48 | 0.33 | 0.38 | 0.14 | -0.46 |

\* MAF: Major allele frequency; Ho: observed heterozygosity; $H_S$ : heterozygosity observed within subdivided populations; $H_T$ : heterozygosity observed across total populations; $F_{ST}$ : Fixation index; $F_{IS}$ :

Based on the allele information at the 33 loci of the 719 lesions, a total of 380 multilocus genotypes (MLG) were identified. Among them, 350 (>92.2%) MLGs were unique to single lesions; MLG359 was the most predominant genotype, 110 lesions of 16 isolates collected from 13 cultivars growing in CA and AZ in 2016 had this genotype, including 3 lesions of UA201614C (Pfs17) and 2 lesions of UA201627D (UA201621A type). MLG 345 consisted of 26 lesions from four isolates in AZ, CT, MA, and WI during 2010, 2016, and 2017 including 18 lesions of UA0510C (Pfs13), 2 of UA201650 (Pfs14), 2 of UA201741B (UA201741B type), and UA201741C (UA201720B type). Other 28 MLGs consisted of 2 to 10 lesions. The 380 multilocus genotypes could be differentiated with the 33 SNP markers.

Using unsupervised discriminant analysis of principal components [41], the 719 lesions were split into four groups (subpopulations), with 208, 283, 151, and 77 lesions respectively (Fig. 2). Most samples could be classified into one group with high probabilities. Among the 719 samples, 662 (92%) had very high posterior probabilities with greater than 0.90; 14 greater than 0.80 belonging to one group. Two lesions had about equal probabilities to one subpopulation or the other.

**Fig. 2.**
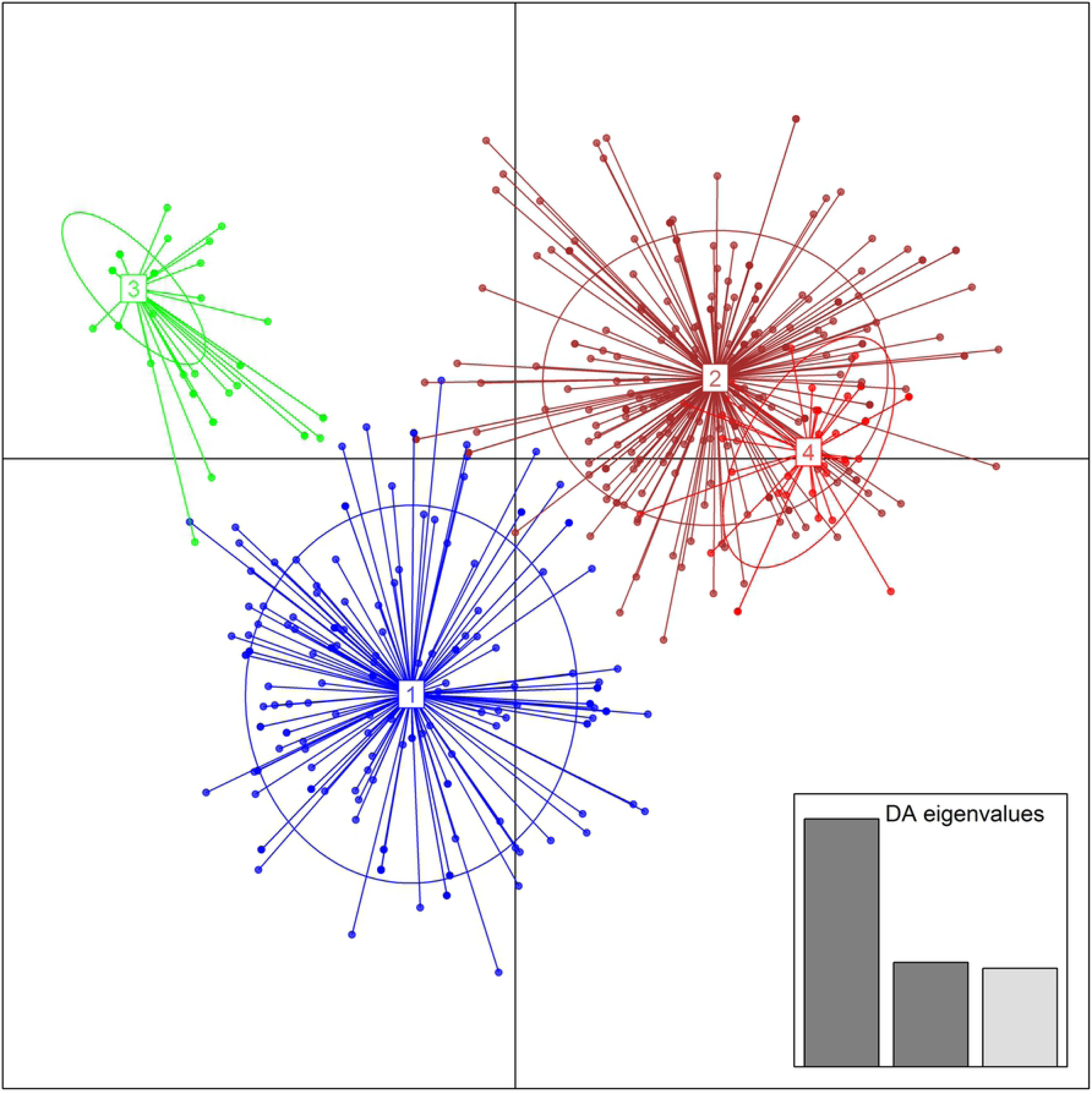
Scatterplots of the DAPC of 719 lesions of spinach downy mildew isolates. This scatterplot shows the first two principal components of the DAPC of lesions, using lesions ID as prior clusters. Groups are shown by different colors and inertia ellipses, while dots represent individual lesions.

### Genetic diversity within spinach downy mildew composite isolates

Nine isolates (one each from NV and MA, two each from CA and MO, and three from ME) had sequence data from two to 13 lesions, but only one genotype was found within each isolate, no variation was found within each of these isolates, indicating clonal reproduction. Five isolates (UA201611G, UA201615D, UA201615H, UA201812B, and UA201818B) had more than two lesions, but only had two MLGs within each isolate (for example, Fig. 3a). Isolate UA201615G had 24 lesions: 4 belonged to MLG1; 8 to MLG2; and 12 to MLG359, only limited variation was found within these field isolate samples. However, there were much more variations observed from other samples. For examples, the isolate UA201611H had 31 MLGs from 37 lesions, and UA201611I had 38 MLGs from 39 lesion of the isolate (Table 1). Those lesions were usually separated into two clades (for example, Fig. 3b).

**Fig. 3.**
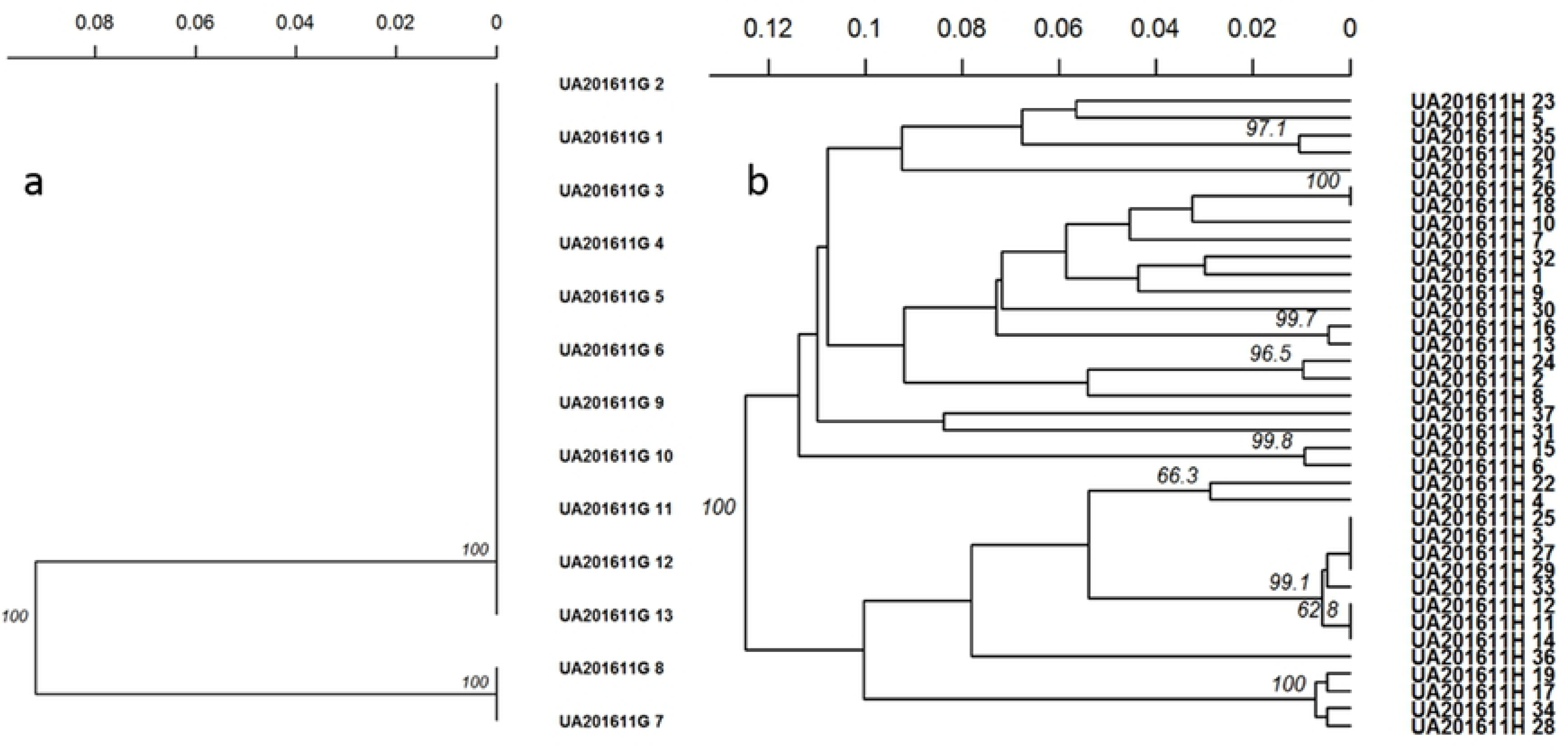
UPGMA dendrograms showing clusters of lesions within one sample based on Nei’s distance. a. cluster analysis of 13 lesions of isolate UA201611G; b. cluster analysis of 37 lesions of isolate UA201611H

The lesions from 6, 25, 6, and 4 isolates were solely classified into Groups 1, 2, 3 and 4 respectively. Six isolates each had samples belonging to Groups 1 and 2, Groups 1 and 3, and Groups 2 and 3. Four isolates had samples belonging Groups 2 and 4. The lesions from four isolates were separated into Groups 1, 2, and 3.

### Genetic diversity among spinach downy mildew isolates

The analysis of molecular variance of the SNP data with or without clone correction indicated that there were significant differences among the 67 isolates of spinach downy mildew pathogen.

The diversity parameters of each isolate after clone correction were listed in Table 1. Wide ranges were found in Shannon-Wiener Index (H) and Simpson’s Index (λ), indicating no variation to abundant variation within different isolates. Lesions from each of the 47 isolates with more than two genotypes were subjected to the index of association analysis (I_A_ and 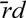) with null hypothesis that alleles recombined freely into new genotypes during the process of sexual reproduction. The results indicated that no evidence to reject the null hypothesis (p>0.05) in two isolates, UA201611L and UA201641A with or without clone correction and in three additional isolates, UA201631, UA201648, and UA201812A after clone correction, suggesting that sexual reproduction existed in these isolates. The null hypothesis was rejected for other isolates (P<0.05), such as UA201615G and UA201611I, suggesting clonal reproduction in those isolates.

The genetic distances between each pair of the 67 isolates ranged from 0.00 to 0.52, within average of 0.17. The large genetic distances with >0.50 were observed between UA201821B and UA201815B, and between UA201648 and UA201615F, while small genetic distances were observed between pair of UA201611E, UA201611F, UA201615A, UA201615D, and UA201645. The isolate UA201615E had highest genetic distance greater than 0.40 paired with other five isolates, averaged 0.293 with all other isolates.

Cluster analysis also showed that isolate UA201615E was an outlier from other isolates (Fig. 4). Isolates UA201648, UA201732, UA201741A, and UA201615F clustered in one distinct clade. Interestingly, some isolates from the same race did not cluster together. For example, three Pfs12 isolates UA201648, UA201630, and UA201718C, four Pfs17 isolates US201608C, UA201614C, UA201718, and UA1014APLP, were placed in different clades, which indicated that clades based on Nei’s distance did not match race classifications (Fig. 4).

**Fig. 4.**
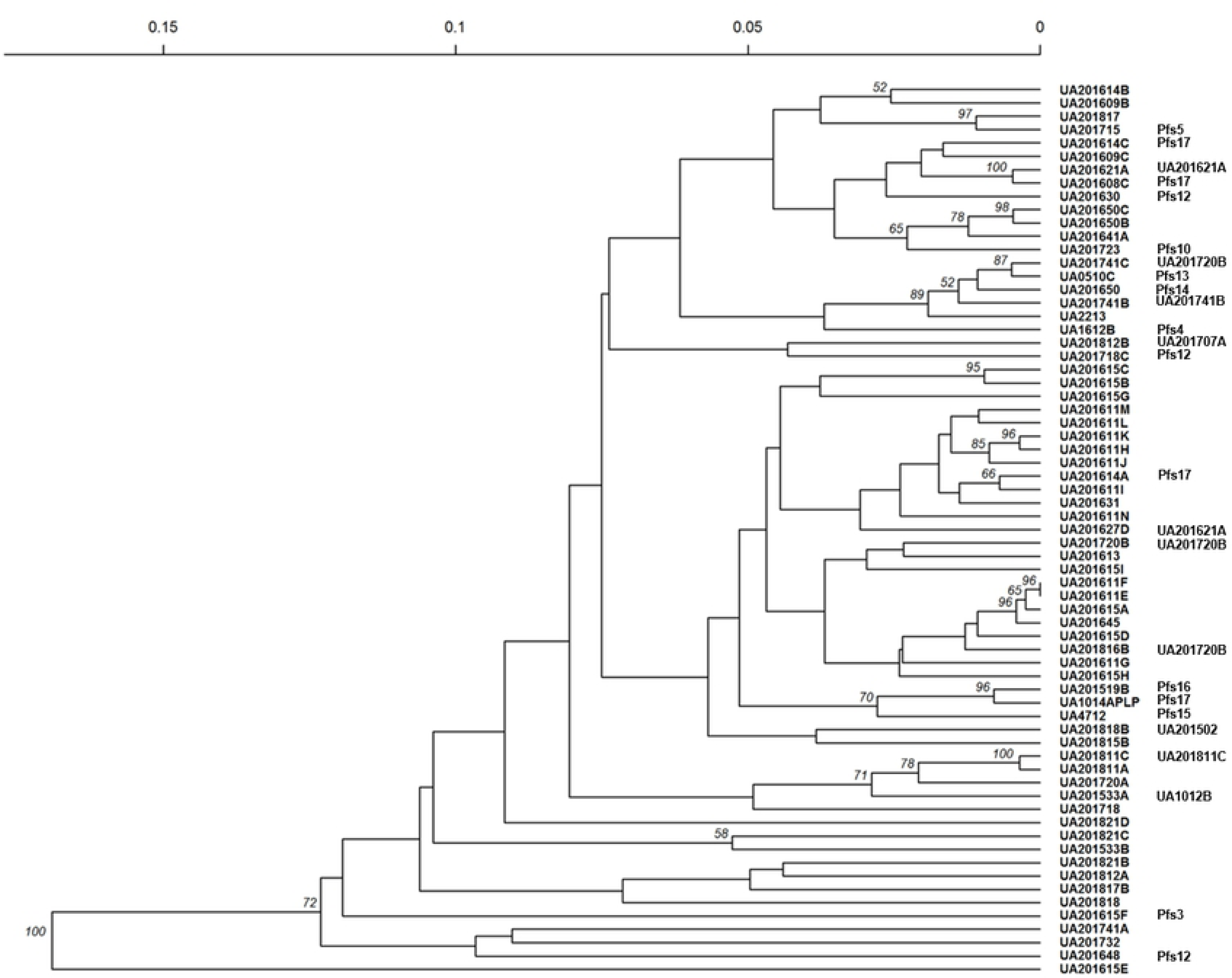
UPGMA dendrogram of the lesions of spinach downy mildew isolates based on Nei’s distance. Bootstrap repeated 1000 times, and bootstrap values greater than 50% were labelled at nodes.

### Host cultivar impact on the genetic diversity of spinach downy mildew isolates

Out of the 67 isolates, three were collected from unknown cultivars; 40 collected from 40 different cultivars, and 2 to 5 from 9 cultivars. In 2016, five isolates UA201611F, UA201611H, UA201611J, UA201615C, and UA201615G were collected on cultivar Platypus. Out of the 91 lesions from these five isolates, 69 belonged to Group 1, 21 belonged to Group 3, and 1 belonged to Group 2. Four lesions of isolate UA201611F, 2 out of the 11 lesions of UA201615C, and 12 out of the 24 lesions of UA201615G, had the same genotypes (the most prevalent genotype MLG359). Four isolates were collected on cultivar Corvair, All lesions of isolate UA201648 (18 total) and UA201818 (5 total) belonged to Group 2; five lesions of isolate UA201650 belonged to Group 4, and one belonged to Group 2, while isolate UA201718C had five Group 2, and one Group 1 lesions. Three isolates, UA201611L, UA201718 and US201611G, were collected on cultivar Sioux, each of the first two isolates had 6 lesions of Group 1 or 2, UA201611G had 2 lesions of MLG 134 (Group 2), and 11of MLG359 (Group 3). The original UA0510C isolate was collected on cultivar Charger, all the lesions of UA0510C belonged to Group4, amongst 18 had the same genotype MLG349.

### Spatial variation among spinach downy mildew isolates

The 67 spinach downy mildew isolates were collected from 13 states. From the two main fresh spinach production states, 403 lesions of 31 isolates collected from CA were sequenced, and 213 genotypes were identified; 175 lesions of 13 isolates collected from AZ were sequenced, and 113 genotypes were identified (Table 3). One to four isolates were collected from other 11 states, and 4 to 36 lesions were sequenced, and 2 to 10 genotypes were identified from each of these states (Table 3).

**Table 3.**
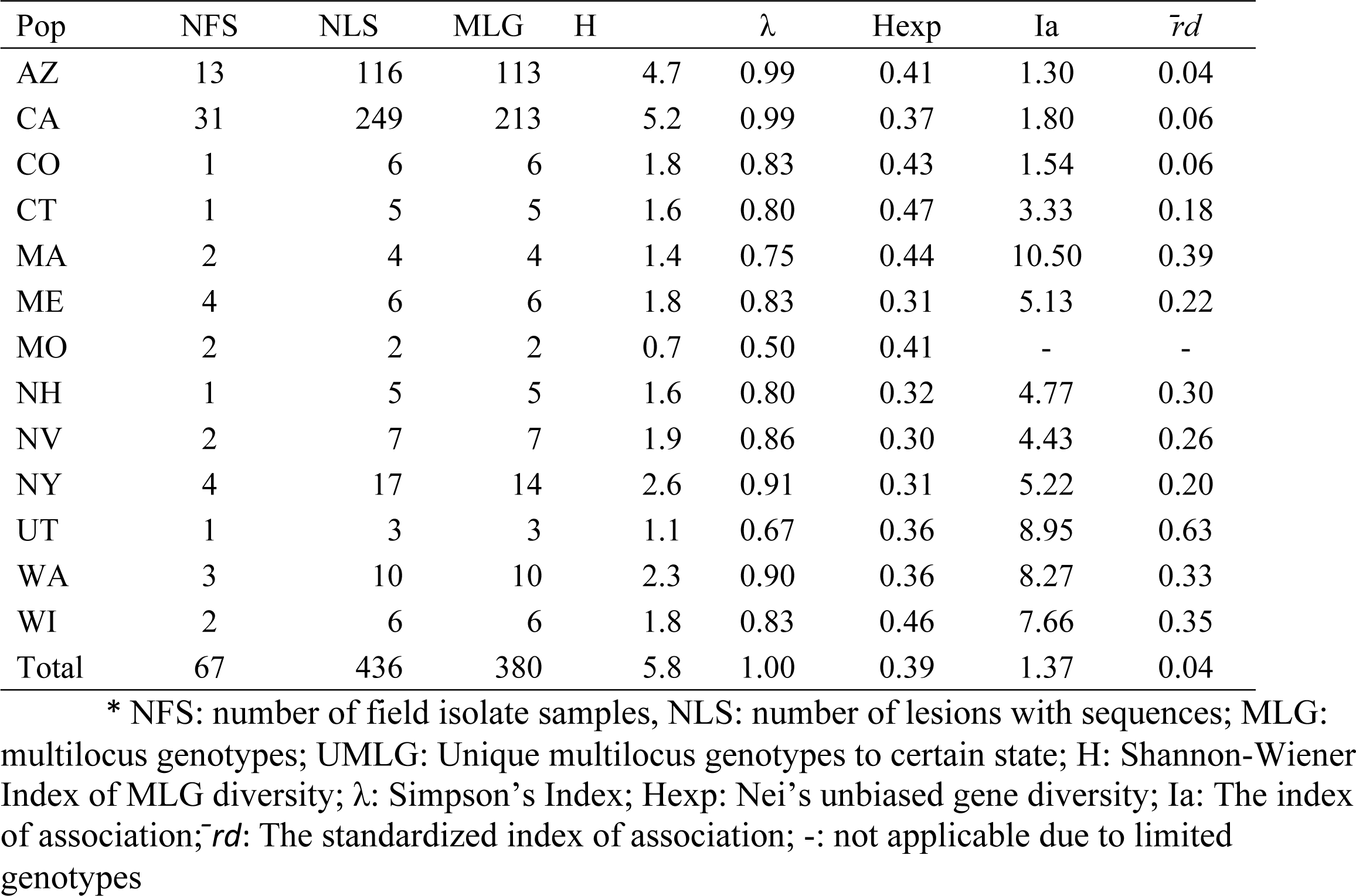
Summary of spinach downy mildew isolates collected from 14 states and diversity parameters after clone correction*

The analysis of molecular variance indicated that there was a significant difference among the *P. effusa* populations collected from different states (P<0.05). The highest Shannon-Wiener Indices (H) were found in the populations of CA and AZ, and the lowest index from the MO population. The Simpson’s Index showed similar results, despite a small range of expected gene diversity (Hexp) from the populations of different states. Evenness (E.5) values indicated equally abundant genotypes in CO and MO populations, while in CA and AZ, the genotypes were not even, some genotypes were more dominant. The index of association analysis (I_A_ and 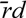) of the populations from 13 states with or without clone correction rejected the null hypothesis at the strata state level (P<0.05) (Table 3). The MO isolates had the largest genetic distances to those from other states, followed by the isolates from ME and NH. Small genetic distances were also found between those from CA and AZ, or from MA and CT, as shown in the dendrogram of cluster analysis (Fig.5).

**Fig. 5.**
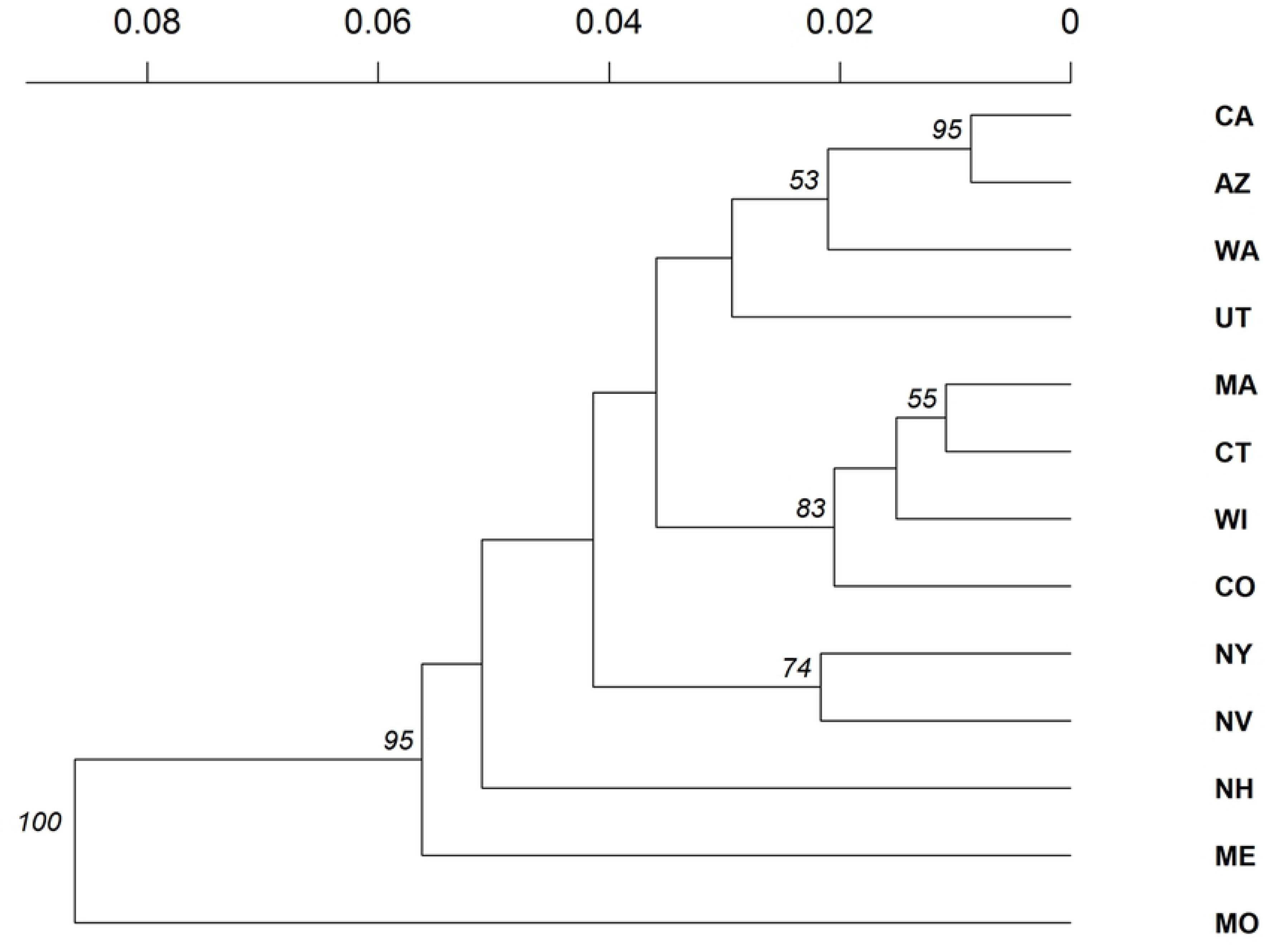
UPGMA dendrogram of the lesions of spinach downy mildew isolates from different states based on Nei genetic distance with 1000 bootstrap replicates. Node value greater than 50% were shown.

There were 369 out of the 380 genotypes specific to only one state, amongst, 209 and 102 genotypes were unique to CA or AZ. Five genotypes were only present in both CA and AZ, including the dominant genotype MLG359, which consisted of 110 lesions. MLG77 and MLG80 contains lesions from WA and UT. MLG349 contained lesions from NY and NV. MLG356 contained 7 lesions from CA, AZ, and NY. MLG345 consisted of 26 lesions from CA, AZ, WI, and CT.

The posterior probabilities of these samples belonging to each group were organized by states (Figure 6). Group 2 is most frequent, which could be found in all states; followed by Group 4, which could be found in 7 states, Group 1 and Group 3 were mainly found in CA and AZ, except one lesion each was found in NH and NY. All four group of lesions were present in the main spinach production states CA and AZ. All lesions from four states, ME, NV, UT, and WA, belonged to Group 2, while all the samples from CO, CT, MA, MO and WI were classified to Groups 2 and 4 (Fig. 6)..

**Fig. 6.**
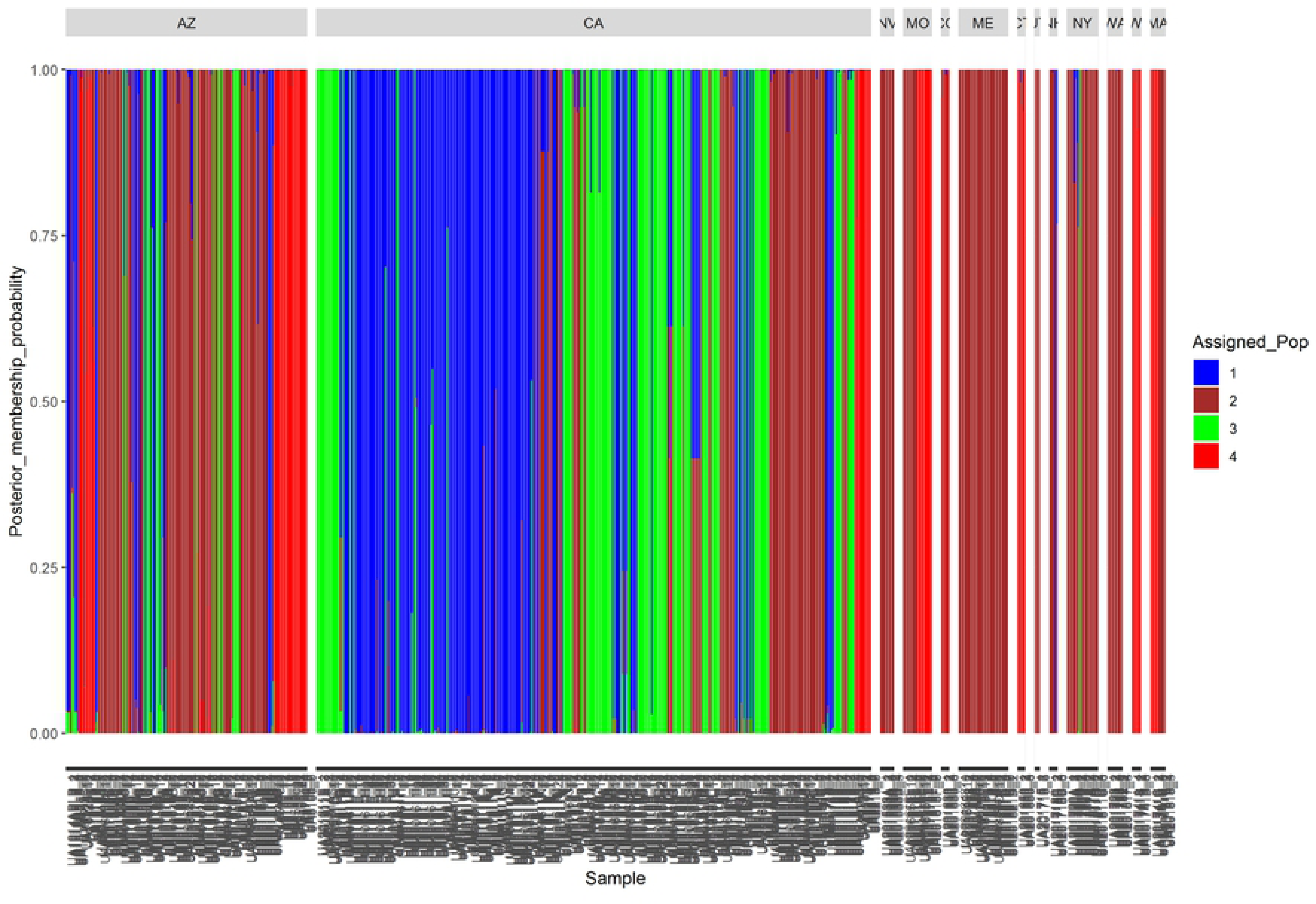
The posterior membership probabilities of the 719 lesion of the 69 spinach downy mildew isolates collected from 13 states.

### Temporal variation among spinach downy mildew isolates

In this study, one to three isolates were collected from 2010 to 2015, but 37, 8 and 15 isolates were collected in 2016, 2017 and 2018 (Table 4). The lesions from each year ranged from 9 to 525, and the number of multilocus genotypes ranged from 3 to 284. The analysis of molecular variance indicated that there was a significant difference among the *P. effusa* populations collected from different years (P<0.05). The highest Shannon-Wiener Index of MLG diversity (H) and Taylor’s Index of MLG diversity (λ) were found in 2016. The index of association analysis (Ia and 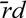) of the SNP data with or without clone correction rejected the null hypothesis of free mating at the strata level of year.

**Table 4.** Summary of spinach downy mildew isolates collected from 8 years and diversity parameters after clone correction

| Pop | NFS | NLS | MLG | H | $\lambda$ | Hexp | Ia | $\bar{r}d$ |
| --- | --- | --- | --- | --- | --- | --- | --- | --- |
| 2010 | 1 | 24 | 6 | 1.79 | 0.83 | 0.41 | 4.15 | 0.38 |
| 2012 | 2 | 13 | 13 | 2.56 | 0.92 | 0.40 | 1.04 | 0.05 |
| 2013 | 1 | 12 | 3 | 1.10 | 0.67 | 0.44 | 8.25 | 0.75 |
| 2014 | 1 | 9 | 5 | 1.61 | 0.80 | 0.30 | 1.49 | 0.09 |
| 2015 | 3 | 12 | 9 | 2.20 | 0.89 | 0.31 | 4.13 | 0.18 |
| 2016 | 36 | 525 | 284 | 5.52 | 0.99 | 0.39 | 1.63 | 0.05 |
| 2017 | 8 | 48 | 41 | 3.70 | 0.98 | 0.39 | 1.49 | 0.05 |
| 2018 | 15 | 76 | 28 | 3.31 | 0.96 | 0.34 | 2.00 | 0.07 |
| Total | 67 | 719 | 380 | 5.83 | 1.00 | 0.39 | 1.37 | 0.04 |

NFS: number of field isolate samples, NLS: number of lesions with sequences; MLG: multilocus genotypes; UMLG: H: Shannon-Wiener Index of MLG diversity; $\lambda$ : Simpson's Index; Hexp: Nei's unbiased gene diversity; Ia: The index of association; $\bar{r}d$ : The standardized index of association.

Among the 380 MLGs identified, 373 MLGs were unique to the year the isolates were collected. Only seven genotypes were present in two or three years, including three genotypes were only present in 2017 and 2018, MLG66 was found in 2014 and 2015, MLG349 was found in 2015 and 2018. Two MLGs were presented in three years, included MLG 345 was presented in 2010, 2016, and 2017; MLG356 was presented in 2016, 2017 and 2018. The smallest genetic distances were found between 2010 and 2013 populations, and between 2017 and 2018 populations, followed by the genetic distances among 2016 and 2017, 2018 populations. Cluster analysis showed that 2010, 2012 and 2013 samples were split in one clade, while others were in a different clade (Fig. 7).

**Fig. 7.**
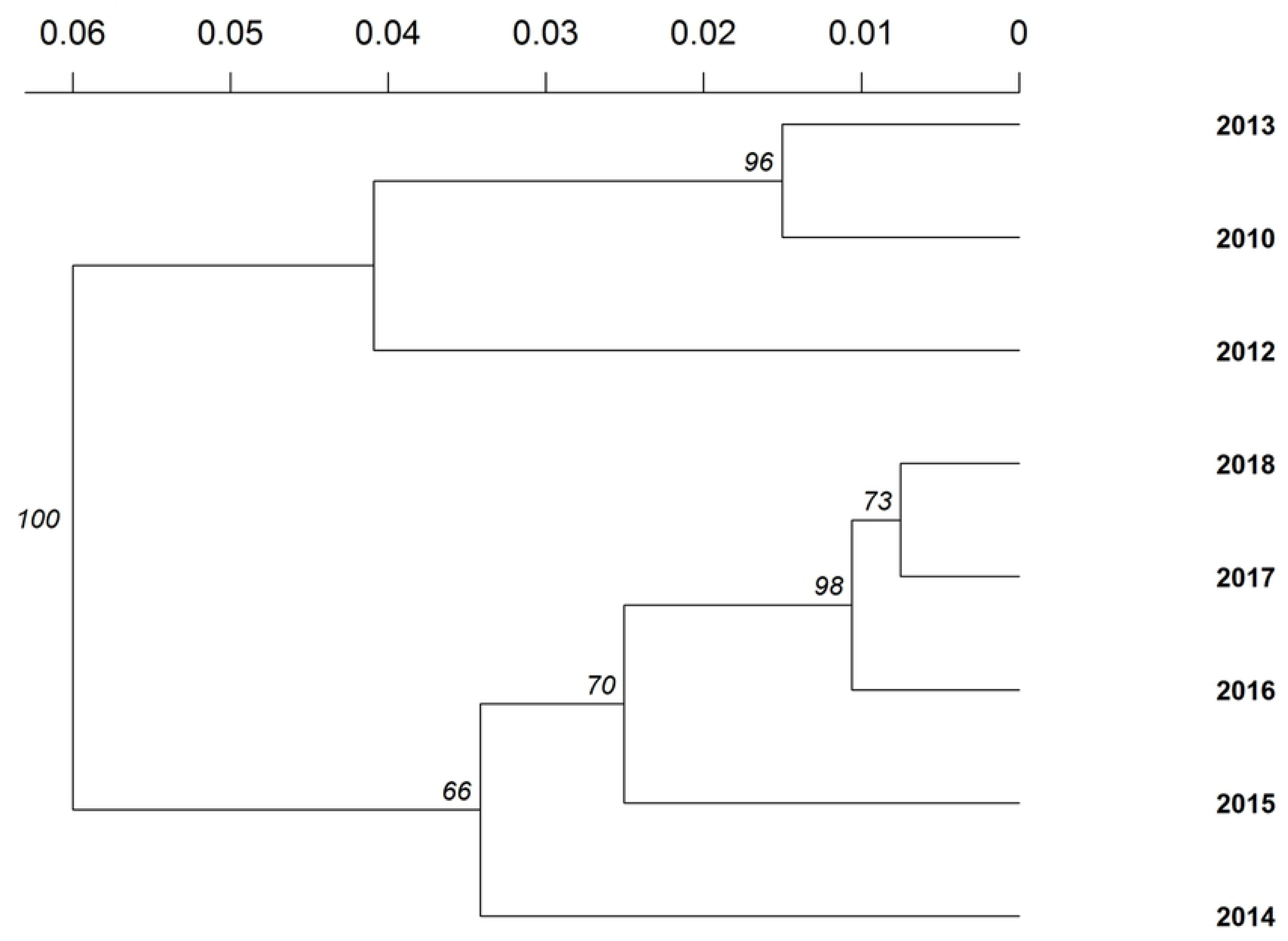
UPGMA dendrogram of the lesions of spinach downy mildew isolates from different states based on Nei’s genetic distance with 1000 bootstrap replicates. Node value greater than 50% were shown.

The posterior membership probabilities of the lesions collected from different years were shown in Figure 8. All lesions of 2010 and 2013 belonged to Group 4, and samples from 2012 belonged to Group 1 and Group 4. Group 2 samples emerged since 2014, Group 3 samples emerged since 2016. All four groups of lesions were found in 2016, 2017 and 2018 (Fig.8).

**Fig. 8.**
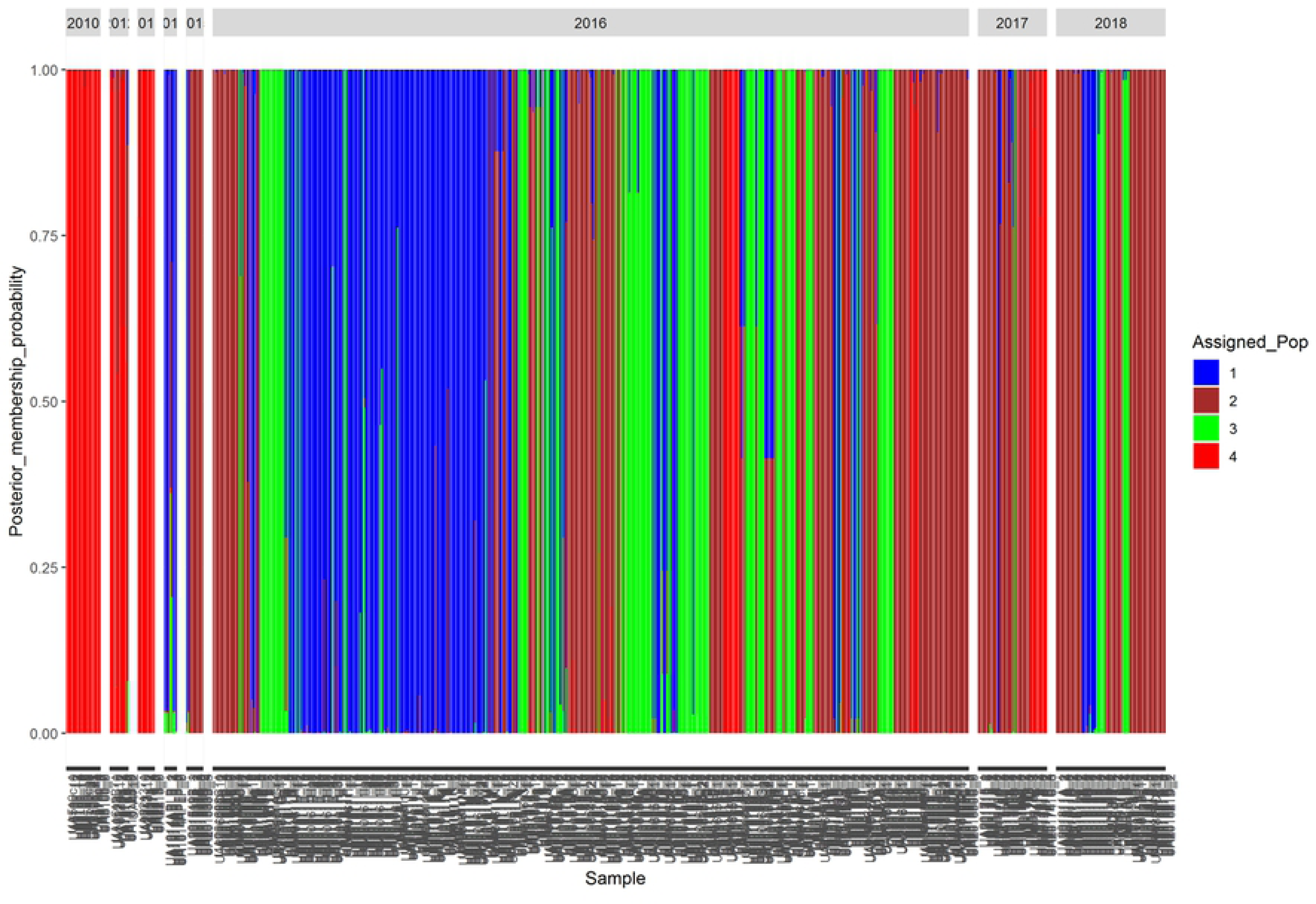
The posterior membership probabilities of the 719 lesions of the 67 spinach downy mildew isolates collected from different years.

## Discussion

Downy mildew of spinach remains a major economic constraint for both organic and conventional spinach production in all major spinach growing areas globally. Although virulence diversity of the downy mildew pathogen, *P. effusa*, has been examined extensively[4–8], little is known of the genotypic diversity of the pathogen at various levels of hierarchy. The genotypic diversity in populations of *P. effusa* has been evaluated with samples collected from CA and AZ [10]. A total of 67 isolates collected from 13 states between 2010 and 2018 was used in the current study, and the genetic diversity was evaluated within and among isolates based on the impact of host cultivar, origin, and year of collection. It is expected to get a better idea about the population structure when samples collected from more states were used. We also recognized that the isolates were opportunistic, and samples for other spinach production states, and other countries, and samples collected in more years need to be used in the future studies.

Considerable variation in the *P. effusa* population was detected by the genotypes of 719 lesions of 67 isolates at the 33 SNP loci examined. A total of 380 multi-locus genotypes were found from the 719 lesions and 357 multilocus genotypes (MLG) were unique to a single lesion. One genotype (MLG 359) was the most predominant genotype, which included 110 lesions representing 16 isolates collected from 13 cultivars from CA and AZ in 2016. A second MLG (MLG345) contained 26 lesions from four isolates collected from four states in 2010, 2016, and 2017. Previous research also found up to 27 isolates had one genotype [10]. It is unknown if this is due to clonality so that some genotypes are much more abundant while others are very rare. Higher resolution may be achieved by using more SNPs or a whole genome sequencing approach. Some genome resources of *P. effusa* are now available and a large number of SNPs have been identified [18, 42]. It may be possible to get additional insight of the population structure and evolution of *P. effusa* by using genome-wide SNPs.

One main difference between oomycetes and fungi is that fungi are haploid while oomycetes are diploids or polyploidy during their life cycle [17, 43]. In this study, only two alleles were found for each of the 33 loci that had been sequenced, indicating the *P. effuse* is a diploid which is consistent with the results from previous research [10]. However targeted sequencing or whole genome sequencing with more isolates of *P. effusa* is needed to reveal the variation in ploidy of *P. effusa*.

A common assumption in population genetics is that diversity is driven primarily by random mating (sexual reproduction), and other forces such as mutation and selection play a reduced or minimal roles as drivers of genetic population diversity. These assumptions are not fully applicable to fungi and oomycetes due to high levels of clonal reproduction. Thus, tools to analyze population structure and genetic diversity developed under this assumption, for example STRUCTURE [44, 45], may not be the best choice for analyzing the genetic diversity and population structure of oomycetes such as *P. effusa* [10]. Recently, the R package *poppr* was developed to facilitate analyses of populations that are clonal or mixed reproduction [16, 36, 37] and has been used for population studies on oomycetes such as *Phytophthora ramorum* [46] and *Plasmopara viticola* [16]. Using poppr and several other R packages, the spatial and temporal impacts on the genetic diversity of *P. effusa* as well as linkage equilibrium were evaluated in this study.

The influence of sexual reproduction on the population structure of *P. effusa* remains unknown. In this study, the index of association analysis (I_A_ and 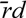) indicated that sexual reproduction existed in a few isolates, and a high level of variation was observed among these isolates. Oospores of *P. effusa* have been observed from spinach seed [12–14] and spinach plants in one downy mildew sample of 2008 [5], two samples in 2016, and 28 samples in 2018 collected from CA (Correll et al., unpublished). Difference was found in the mating proteins in the genome sequences of isolates of *P. effusa* [42], oospores can be produced by co-inoculation of two isolates selected accordingly to different mating features [47], indicating heterothallic sexual reproduction occurs in *P. effusa*, which was also found in several other oomycetes, such as *Phytophthora*, *Bremia*, and *Plasmopara* [21]. Among those isolates that rejected sexual reproduction hypotheses, nine isolates contained multiple lesions, but only one genotype was observed, suggesting clonal production among these isolates of *P. effusa*. There could be large amount of variations in some clonal reproduced isolates, such as the isolates UA201611H and UA201611I, which had 37 and 39 lesions, but 31 and 38 multilocus genotypes were observed. Variation generated by asexual reproduction was also observed in other oomycetes, such as *Phytophthora capsici* [48] and *Plasmopara viticola* [16]. The co-existence of both sexual and asexual reproduction may result in the fast emergence of the new races/novel strains of the pathogen *P. effusa*.

Although there was only one genotype in some composite isolates, other composite isolates typically contained multiple genotypes, and more variations were found among isolates of the same race. Thus, a given race typically represents a population of isolates with one, or a set of common avirulence genes that can interact with a corresponding resistance gene in spinach. Although the develop molecular diagnosis tools for rapid race identification of *P. effusa* would be very valuable, the significant amount of variation observed in this study suggests that this would be exceptionally difficult unless the avirulence genes of the races were identified and the molecular markers were developed targeting the avirulence genes.

Host cultivars could have a significant impact on oomycete pathogens. Oomycetes absorb nutrients from host plants, the obligate pathogens rely more on host plants due to their defects in metabolism [20]. It was observed that the imbalance in ligin biosynthesis in *Arabidopsis thaliana* could promote the sexual reproduction of *Hyaloperonospora arabidopsidis* [49]. The resistance in host cultivars exert selection pressure on the pathogen, which can filter out any strains of the pathogen that cannot overcome the resistance, thus affecting the genetic diversity of the pathogen. For example, the spinach cultivar Lion is susceptible to races Pfs 10 and Pfs 17, but has resistance to races Pfs 1-9, 11-16 [6]. Thus, isolates of races Pfs10 or Pfs 17 may be recovered from cultivar the cultivar Lion, but it is not possible to recover isolates of other named races of *P. effusa*.

In addition to host cultivar, spatial and temporal variation also was found in the population of *P. effusa*. When a race of *P. effusa* occurs in a spinach production area, cultivars with resistance to this race will be deployed until the resistance is broken down by new races. While the battle is continuous, spinach cultivars with different combinations of resistance genes are grown in different regions at different times. This aspect of production likely explains the significant spatial and temporal effects on the variation of the pathogen. However, it is unknown if there are any environmental (temperature, humidity, etc.) effects on the diversity of the pathogen population.

Overall, a great deal regarding the genetic diversity of populations of *P. effusa* has been learned from this research. The genetic diversity of *P. effusa* population may be affected by either or co-exsitance of sexual and asexuaul reproductions, host selection pressure, in addition to spatial and temporal effects. The fast emergence of new races may be driven by multiple molecular mechanisms. The ability to do controlled crosses and generate sexual progeny of *P. effusa* represents a significant advance in our understanding of genetic diversity of this important pathogen [47]. The relative importance of asexual variation versus sexual variation in virulence in the *P. effusa* population will help determine the rate at which novel strains overcoming the deployed resistance can develop.

